# The copine protein NRA-1 regulates activity-dependent cholinergic signalling and sensory integration in *C. elegans*

**DOI:** 10.64898/2026.07.09.737470

**Authors:** Leona Cesar, Jana F. Liewald, Guillermina Hernando, Emelie Aspholm, Alexander Kolsrud, Sebastian Wabnig, Cecilia Bouzat, Alexander Gottschalk, Julia Morud

**Author notes:** Equal contribution (Leona Cesar, Jana F. Liewald).

## Abstract

Membrane-associated proteins regulate the localisation and function of ligand-gated ion channels, yet how they shape synaptic efficacy and behaviour remains poorly understood. Here, we identify the copine family protein NRA-1 as an activity-dependent regulator of cholinergic signalling and sensory circuit function. Electrophysiological recordings at the neuromuscular junction of *C. elegans* revealed that loss of *nra-1* does not alter responses to acute agonist application or the initial response to synaptic stimulation but selectively impairs sustained and repetitive cholinergic transmission during ongoing activity. Single-channel recordings further demonstrated that NRA-1 does not affect the unitary conductance of levamisole-sensitive acetylcholine receptors (L-AChRs) but instead regulates receptor gating by increasing channel closed times and reducing opening frequency, resulting in an overall decrease in receptor activity. Despite these synaptic defects, *nra-1* mutants displayed normal baseline locomotion but exhibited impaired chemotaxis, abnormal food localisation and defective egg-laying behaviour. Together, our findings identify NRA-1 as an activity-dependent regulator of postsynaptic receptor function that sustains cholinergic signalling during repeated activity and links receptor dynamics to sensory behaviour. These results establish copine proteins as important modulators of synaptic efficacy and suggest that activity-dependent control of receptor function represents a conserved mechanism for tuning neural circuit performance.

## Introduction

Animals rely on their sensory systems to navigate complex environments, locate food and avoid harmful stimuli. In the roundworm *Caenorhabditis elegans* (*C. elegans*), sensory integration underlies essential behaviours, such as chemotaxis, food seeking, egg-laying and avoidance responses^1–5^. Owing to its genetically tractable nervous system, *C. elegans* is a powerful model system for dissecting neural circuit and molecular mechanisms underlying behaviour^6–8^. Central to neural circuit function are ligand-gated ion channels (LGICs), which mediate fast synaptic transmission by converting chemical signals into rapid changes of neuronal membrane potential, thereby shaping behavioural outputs^9,10^. Chemotaxis provides a robust and quantifiable behavioural readout of sensory function in *C. elegans*, requiring animals to detect, integrate and respond to chemical gradients^11–14^ through coordinated activity and precise transduction of sensory neurons, interneurons and motor neurons^12,15^, including the availability and activity of LGICs that modulate synaptic strength and circuit dynamics downstream of the odour recognition^16,17^.

For the execution of complex behaviours, animals rely on higher-order neuronal circuits that coordinate the detection and integration of sensory cues and interact with basal motor machinery to generate appropriate behavioural responses^18,19^. Proteins involved in synaptic function may act at multiple levels within these systems, influencing both the basal neuromuscular system as well as higher-order sensory integration circuits, thereby complicating the interpretation of mutant phenotypes. The neuromuscular junction (NMJ) is central to locomotion^20,21^. Premotor interneurons regulate the activity of motor neurons that stimulate, and in some invertebrate systems, inhibit^22^, muscle function that coordinates movement^23^. Motor neurons release neurotransmitters such as acetylcholine, which excite the muscle via nicotinic acetylcholine receptors (nAChRs), pentameric ligand-gated ion channels (pLGICs), on muscle cells. Muscle activation depends on receptor abundance and their functional properties, such as gating and closing kinetics as well as desensitisation^21^. LGICs are dynamically regulated by accessory proteins, in an activity-dependent manner^24^, helping maintain postsynaptic responsiveness under conditions of altered presynaptic input^25^. At the *C. elegans* NMJ, biochemical and proteomic studies identified several proteins associated with the nAChRs, including the copine family protein NRA-1^26^, which are Ca^2+^-dependent membrane-binding proteins^27^. Copines, such as Cpne6, have been implicated in membrane trafficking, spine structural plasticity and neuronal development^28–30^. Loss of NRA-1 reduces nAChR expression at the muscle surface in *C. elegans*, indicating a role for NRA-1 in pLGIC plasma membrane insertion or stabilisation^26^. In addition, silencing of the gene with feeding RNAi, significantly increase both levamisole and nicotine resistance, while mutating *nra-1* cause reduction of nAChRs, but not GABARs^31^. However, despite these insights at the NMJ, the broader roles of NRA-1 and copine proteins in ion channel regulation and neural circuit function remain poorly understood.

Chemotaxis in *C. elegans* depends on specialised chemosensory neurons that detect environmental cues via G protein-coupled receptors (GPCRs) localised to sensory cilia^12,32^. Activation of these receptors initiates intracellular signalling cascades that convert chemical stimuli into neuronal activity^33^, which play central roles in modulating ion channel activity and neuronal excitability^17^. Fast communication within these circuits relies on LGICs, which convert presynaptic neurotransmitter release into postsynaptic responses, enabling accurate sensory processing and behavioural output ^7,34–38^. In parallel, membrane-associated proteins, including SNAREs and calcium-binding adaptor proteins, coordinate neurotransmiaer release, regulate receptor trafficking as well as ion channel localisacon and stability ^39–42^. However, how such regulators influence ion channel function and contribute to chemotactic behaviour remains poorly understood.

Here, we investigate the physiological role of the copine protein NRA-1 by combining electrophysiology, single-channel recordings, optogenetics and behavioural analyses. We show that NRA-1 is not required for acute cholinergic transmission but instead sustains cholinergic signalling during repeated synaptic activity by regulating L-AChR function. In *nra-1* deletion animals a strong defect in chemotactic navigation, food leaving and egg-laying was detected, suggesting a broad impairment of sensory integration. These findings provide further evidence that NRA-1 acts as an activity-dependent regulator of ion channel localisation and function in cholinergic NMJs, while also suggesting a novel role for NRA-1 in modulating sensory signalling in *C. elegans*.

## Materials and methods

### Materials availability

Materials generated in this study, including strains, plasmids and clones, are freely available from the lead contact upon reasonable request. Further information and requests for *C. elegans* strains and plasmids is to be sent to and will be fulfilled by the lead contact Julia Morud Lekholm

### C. elegans

Unless otherwise specified, *C. elegans* were maintained at 20 °C on nematode growth medium agar (NGM-Agar) plates seeded with *E. coli* OP50 as a food source^43^. Transgenic rescue lines were generated by microinjecting plasmid DNA into the gonads of young adult hermaphrodites. Progeny carrying stable extrachromosomal arrays were selected and propagated as independent strains. For optogenetic stimulation, animals were raised in the presence of all-*trans* retinal, as previously described^44^. Strains used were: N2 wild type (Bristol strain), RB1068: *nra-1(ok1025)*, ZX460: *zxIs6[punc-17::ChR2(H134R)::YFP; lin-15^+^],* ZX1006: *nra-1(ok1025); zxIs6[punc-17::ChR2(H134R)::YFP; lin-15^+^], JML88: nra-1(ok1025) JemEx03[(podr-10::nra-1 cDNA::SL2 mKate gpd-2 3’UTR); punc-122::GFP],JML89: nra-1(ok1025) JemEx04[(pstr-1::nra-1 cDNA::SL2 mKate gpd-2 3’UTR); punc-122::GFP], JML90: nra-1(ok1025) JemEx05[(pstr-2::nra-1 cDNA::SL2 mKate gpd-2 3’UTR); punc-122::GFP], JML101: nra-1 (ok1025) JemEx11 [pnra-1:nra-1cDNA_SL2 mKate, ccGFP]*.

### Isolation and culture of *C. elegans* muscle cells

To investigate the molecular mechanisms underlying NRA-1 function, electrophysiological recordings were performed on native L-AChRs expressed in L1 muscle cells from both wild type and *nra-1(ok1025) C. elegans*. L1 muscle cells were obtained from *C. elegans* eggs and cultured using a previously established protocol^45^, which has been routinely employed in our previous studies. Electrophysiological experiments were conducted 1–5 days after cell isolation^46–51^.

### Electrophysiology

#### Patch-clamp in C. elegans body wall muscles

Recordings from dissected *C. elegans* body wall muscle were essentially performed as described previously^20,44,52^. After dissection, muscle cells were treated for 12 s with 0.5 mg/ml collagenase (C5138, Sigma-Aldrich, Germany) in extracellular *C. elegans* Ringer’s solution (CRG; 150 mM NaCl, 5 mM KCl, 5 or 0 mM CaCl_2_, as indicated, 1 mM MgCl2, 10 mM glucose, 15 mM HEPES (pH 7.35), ∼340 mOsm) and washed with CRG. The pipette solution contained 120 mM KCl, 20 mM KOH, 4 mM MgCl_2_, 5 mM TRIS-HCl, pH 7.2, 0.25 mM CaCl_2_, 4 mM ATP, 36 mM sucrose, 5 mM EGTA, ∼315 mOsm. We used an EPC10 amplifier with head stage and Pulse software (HEKA), to clamp the cells to -60 mV holding potential. Agonists were pressure applied (Parker Picospritzer III). Light activation was performed using an LED lamp (KSL-70, Rapp OptoElectronic, Germany) at a wavelength of 470nm (8 mW / mm2) and controlled by the HEKA software.

#### Patch-clamp recordings in cultured C. elegans muscle cells

Single-channel currents were recorded from L1 muscle cells using the cell-attached patch-clamp configuration, following established protocols^47,48^. Recordings were performed at 20 °C with a pipette potential of 100 mV to assess the activity of native receptors. ACh or levamisole (Sigma-Aldrich) was included in the pipette solution. Bath and pipette solutions were prepared as previously described^47,48,51^. Channel activity was acquired using the Acquire software and analysed with TAC (Bruxton Corporation). Open- and closed-time distributions, as well as burst duration histograms, were plotted using a logarithmic time axis and square-root ordinate and fitted with sums of exponential components by maximum-likelihood methods using TACFit (Bruxton Corporation).

### Worm tracking

Baseline locomotion was tested in N2 and *nra-1(ok1025)* animals, both strains were maintained under identical culture conditions. Synchronized L4 larvae were transferred to seeded fresh nematode growth medium (NGM) agar plates (18 g of agar, 2.5 g of Bacto peptone and 3 g of NaCl in 1 litre of ddH₂O, autoclaved and then supplemented with 5 ml of KPO₄ buffer, 1 ml of 1 M CaCl₂, 1 ml of 1 M MgSO₄ and 1 ml of 5 mg/ml cholesterol in 96% ethanol) using M9 buffer (3 g of KH₂PO₄, 6 g of Na₂HPO₄, 5 g of NaCl and 1 ml of 1 M MgSO₄ in 1 l ddH₂O, autoclaved) and allowed to develop into day 1 adults overnight. On the day of analysis, animals were transferred using an eyelash pick to unseeded NGM assay plates to minimize handling-induced behavioural perturbations.

13-16 worms were placed within a 1 cm diameter copper ring positioned on each assay plate to restrict movement outside the imaging area. Following transfer, animals were allowed to acclimate for 5 min before behavioural recording. A total of approximately 150 animals per genotype were analysed across multiple independent assay plates and experimental days. Locomotion was recorded with DinoLite 2.0 software for 10 min using a Dino-Lite AM4115-FIT microscope (DinoLite, AnMo Electronics Corporation, Taiwan) equipped with 850 nm infrared illumination and diffuse transmitted backlighting. Videos were analysed using Tierpsy Tracker^53^, which automatically segmented individual animals, extracted body posture information and generated quantitative locomotion features. Mean locomotion speed was calculated for each animal and used for subsequent comparisons between mutants and wild-type controls.

### Chemotaxis

Chemotaxis assays were performed as previously described^54^. Day-1 adult worms were washed from NGM-agar plates before being transferred into 1.5 ml tubes. The worms were allowed to settle by gravity for 5 minutes and then washed four times in M9 buffer to remove residual bacteria. The worms were then transferred to chemotaxis (CTX) plates (20 g agar in 1 l ddH₂O; autoclaved, then supplemented with 5 ml KPO₄ buffer, 1 ml 1 M CaCl₂ and 1 ml 1 M MgSO₄) prepared four days in advance and dried under a sterile laminar flow hood to ensure consistent humidity. For the assays, 2 µl of odorant and control solutions were placed on opposite sides of the CTX plates. Diacetyl (Sigma-Aldrich, USA) was diluted to 0.05% (v/v) in 96% ethanol (Solveco, Sweden) and isoamyl alcohol (IAA; Sigma-Aldrich, USA) to 1% (v/v). Each odorant dilution was mixed 1:1 with 1 M sodium azide (Merck, Germany) immediately before use. As a control, 96% ethanol was mixed 1:1 with 1 M sodium azide. Approximately 50–70 worms were transferred to the centre of each plate in a small drop of M9 using a glass Pasteur pipette and excess liquid was removed with filter paper. Plates were kept covered in a fume hood during the assay and worms were allowed to migrate freely for 60 min at room temperature. The choice index (CI = (n_test_ -n_control_)/n_total_) was calculated as previously described^17,54^.

### 2-nonanone food leaving

The food-leaving avoidance assay was adapted from a previous publication^15^. NGM-agar plates were seeded with 20 µl of *E. coli* OP50, which was prepared as a fresh morning culture from an overnight starter and diluted in LB before being regrown to an OD₆₀₀ of 1. The plates were then incubated at room temperature for 1 hour before the assay was started. Fifteen day-1 adult hermaphrodite *C. elegans* were transferred to the bacterial lawn and allowed to acclimatise for 1 hour. Afterwards, 1 µl of 100% 2-nonanone (Sigma-Aldrich, USA) was applied directly to the agar adjacent to the bacterial patch. Worm behaviour was filmed for one hour using Dino-Lite cameras (model AMZ73115MZTL; AnMo Electronics Corp., Taiwan). Four cameras were used simultaneously to record four plates in parallel. Each genotype was assayed in triplicate per day. Worm positions were analysed from the videos every 5 minutes for the first 25 minutes and then every 10 minutes. Avoidance was quantified as the percentage of worms that had left the OP50 lawn at each time point, using the number of worms initially present on the lawn (time point 0) as the 100% reference value.

### Spontaneous food-leaving assay

The assays were performed on NGM plates, prepared as described above and seeded with 20 µl of *E. coli* OP50 at an OD₆₀₀ of 1. Fifteen young adult hermaphrodites were transferred to the bacterial lawn and allowed to acclimatise for one hour. Plates were then filmed for nineteen hours using the four-camera setup as for the 2-nonanone assay. The number of worms on and off the bacterial lawn was counted hourly throughout the recording and food-leaving was quantified as the percentage of worms off the lawn. The number of eggs laid outside the bacterial lawn was also counted after the 19-hour recording.

### Statistics

The experimental data are shown as mean values with standard deviation (mean ± SD) or mean values with standard error of the mean (mean ± SEM). Statistical comparisons were done using a Student’s t-test, a Welch’s test or a Two-way ANOVA. A significance level of p < 0.05 was considered as statistically significant. Tests were performed with SigmaPlot 12.0 (Systat Software, Inc.) or GraphPad Prism 11.0.

## Results

Guided by previous research demonstrating that loss of *nra-1* reduces synaptic L-AChR expression, indicating a functional link between NRA-1 and cholinergic receptor localisation^26^, we sought to determine whether NRA-1 directly regulates AChRs function and whether this role extends to the modulation of sensory processing. We first examined the neuronal expression pattern of *nra-1* to gain a deeper understanding of where in the worm the protein might play a role. To this end, we used the public single-cell transcriptome dataset CeNGEN^55^ for *nra-1* cell-ID. This revealed that *nra-1* is expressed broadly, almost pan-neuronally, with high expression detected in several interneurons (AIB, AVH, SIA, and SIB) as well in several sensory neurons (AWC, URX, PLM and PLN) and cholinergic ventral cord motor neurons (DB1, DB2, VB1 and VB2) (Figure 1). In addition to neuronal expression, *nra-1* is found in muscle mesoderm, such as body wall muscle tissue^56^, which is in line with previously reported *nra-1* expression profiles^26^.

**Figure 1.**
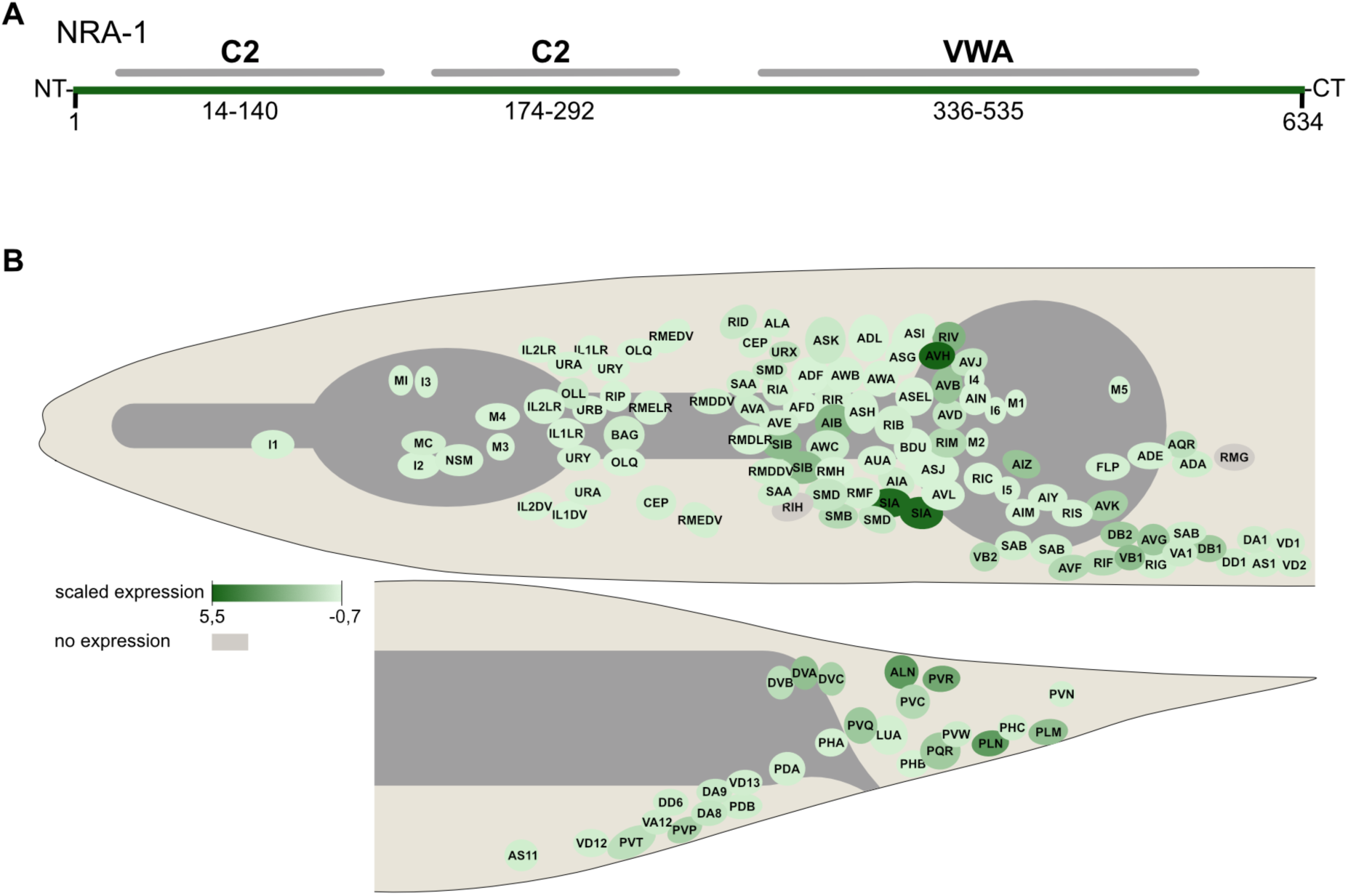
The copine NRA-1 has two C2 domains and broad gene expression in *C. elegans* neurons. A. NRA-1 have two predicted Ca^2+^ binding C2 domains and one von Willebrand factor type A domain (VWA). Functional protein domains predicted using SMART v10^72^. B. *nra-1* gene expression^55^ is found in a large proportion of all *C. elegans* neurons, including cholinergic ventral cord motor neurons (shown only partially for space constraints), interneurons and sensory neurons. In addition, it is also expressed in muscle mesoderm. Expression level is indicated by increasing intensity of green colour. No detected expression is noted in grey. Expression is plotted as scaled expression, equal to transformed raw gene expression for allowing comparisons between genes and cells.

As NRA-1 was initially identified in association with biochemically purified L-AChR, and the *nra-1(ok1025)* mutant was shown to exhibit levamisole and nicotine resistance, along with reduced levels of levamisole receptors at the plasma membrane of muscle cells ^26^, we sought to investigate whether the identified expression of *nra-1* in cholinergic motor neurons might contribute to responses during L-AChR agonist application or intrinsic ACh release in NMJ, possibly in an activity-dependent manner.

To this end, we employed both physiological and acute assays of muscular neurotransmitter receptor sensitivity, together with optogenetic stimulation of cholinergic neurons, to assess the impact of the *nra-1(ok1025)* mutation on ACh release *in vivo*. We first examined general agonist responses at the NMJ, which in *C. elegans* comprises a mixed cholinergic and GABAergic synapse. This synapse includes two types of AChR (the levamisole receptors^57–60^ which are specifically activated by levamisole, L-AChR, and the nicotine-sensitive ACR-16 containing AChR^61^), as well as the UNC-49 GABA_A_^20,62^. We measured currents evoked by puff-applied agonists (0.5 mM ACh, 0.5 mM levamisole, 1 mM nicotine, 0.1 mM GABA; Figure 2). All agonists elicited comparable current responses in muscle cells of wild type animals and *nra-1(ok1025)* mutants. These findings were unexpected, given the reduced abundance of levamisole receptors at the muscle surface of *nra-1* mutants, and since they did not confirm the previously obtained result from paralysis assays^26^.

**Figure 2.**
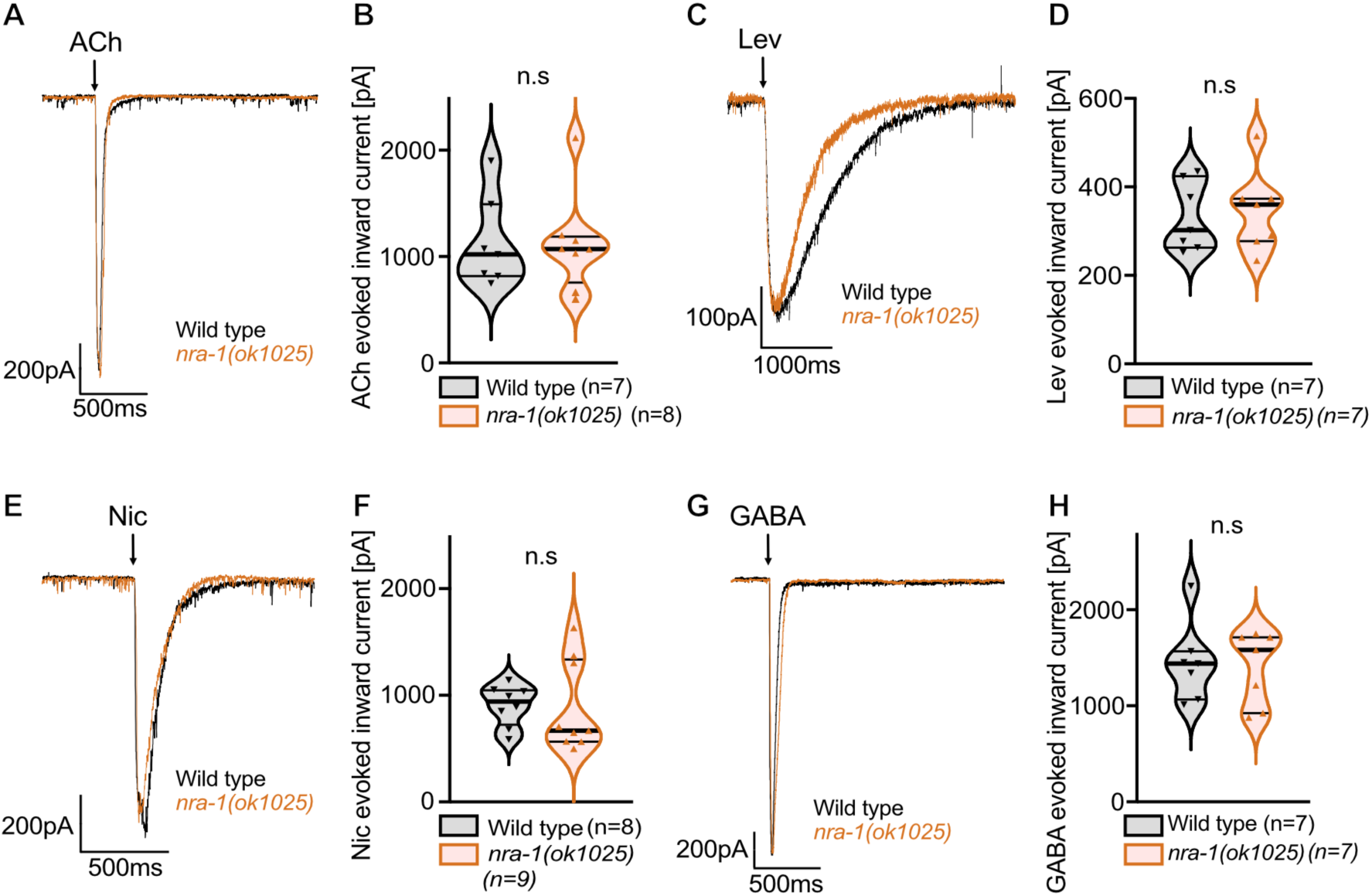
Acute agonist application to the NMJ does not show abnormalities in mutants lacking NRA-1. ACh (0.5 mM) (A-B), levamisole (0.5 mM) (C-D), nicotine (1 mM) (E-F), or GABA (0.1 mM) (G-H) were applied to dissected NMJs, from which postsynaptic currents were recorded. The current traces show representative original records and violin plots show individual measurements as well as median currents with 25-75 % quartiles (thick and thin black lines), min to max, of the indicated number of animals and genotypes. Statistical analyses were performed by unpaired t-test. n.s, no significant difference.

In these agonist application experiments, supraphysiological concentrations of agonist are delivered across the entire muscle cell surface, whereas nAChRs are normally confined to postsynaptic sites. If NRA-1 is required for proper postsynaptic localisation, its absence may not alter overall surface expression levels but instead lead to a redistribution of receptors to extrasynaptic regions. We therefore examined endogenous, spatially restricted ACh release at presynaptic sites, using optogenetic stimulation of cholinergic neurons via channelrhodopsin-2 (ChR2)^44,63^. In response to brief (10 ms) light stimulation, both wild type and *nra-1* mutants exhibited comparable inward currents (Figure 3A), although *nra-1* animals showed a non-significant trend toward reduced currents (mean: 1587 pA in wild type, 1155 pA in *nra-1,* p = 0.12). We next applied longer (1s) light stimuli (Figure 3C), which elicit a rapid peak current followed by sustained ACh release and prolonged muscle activation throughout the duration of the light pulse. While the initial peak responses were similar between genotypes (mean ± SEM; wild type: 1728 ± 241 pA; *nra-1*: 1470 ± 226 pA, p = 0.45), the steady-state current was significantly reduced in *nra-1* mutants (mean ± SEM; wild type: 68 ± 7 pA; *nra-1*: 45 ± 6 pA, p < 0.05).

**Figure 3.**
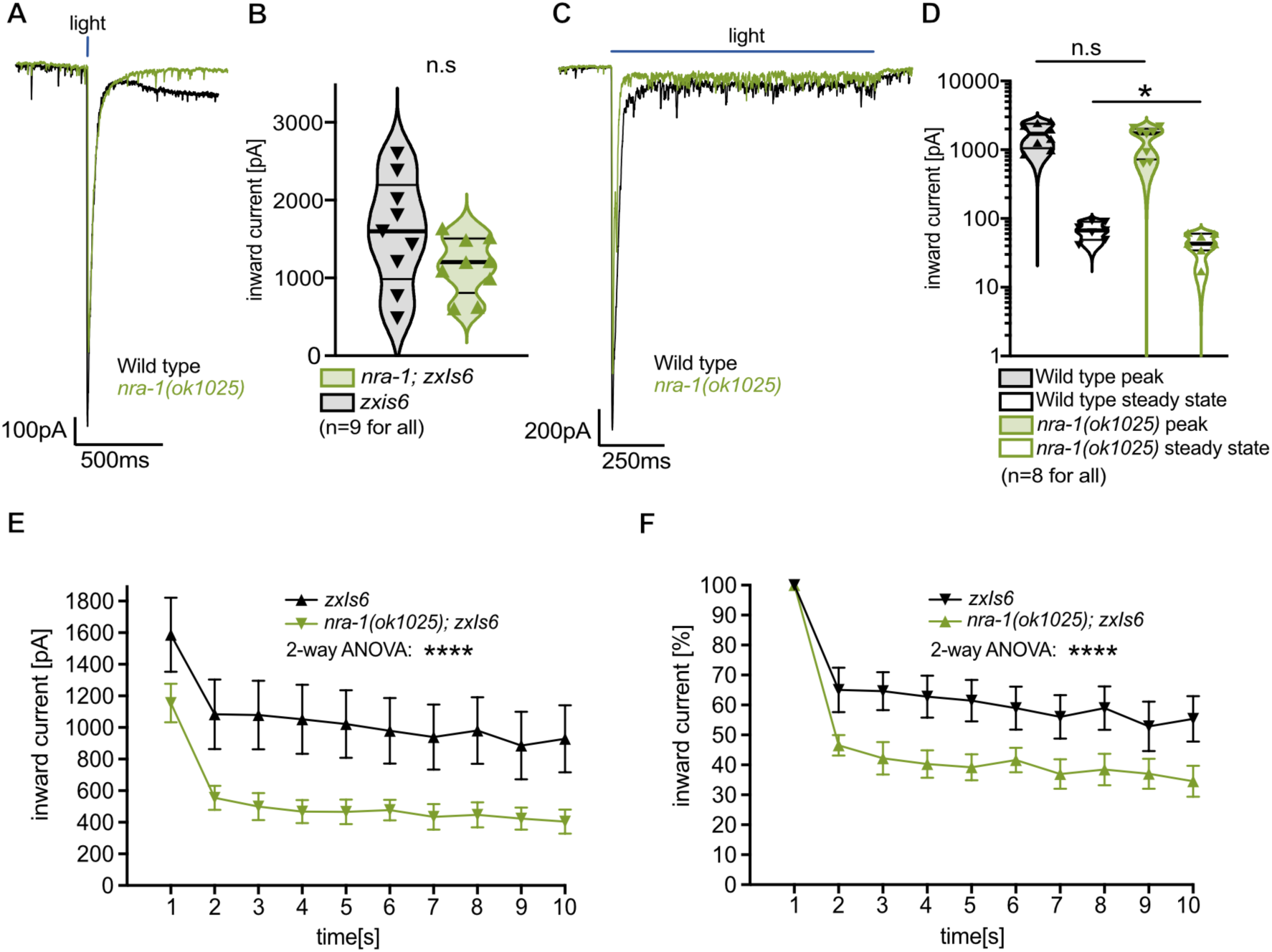
Light-evoked currents in response to brief and sustained optogenetic stimulation of cholinergic motor neurons show reduced currents whereas repeated optogenetic stimulation unravels a strong defect in *nra-1(ok1025)* mutants. Currents in response to brief (10 ms) (A-B) or long (1000 ms) (C-D) light stimulation (470 nm, 8 mW/mm²). The panels show representative original current traces and median currents with 25-75 % quartiles (thick and thin black lines), min to max, of the indicated number of animals and genotypes. E: Repeated optogenetic stimulation is represented as mean ± SEM currents in wild type and *nra-1* mutants, to ten consecutive (0.5 Hz) 10 ms photostimuli (470 nm, 8 mW/mm²). F: Normalised responses, relative to the first response. Statistical analyses were performed by unpaired t-test (note: for the long stimuli, only the initial peaks or the steady state were compared among genotypes) or by 2-way ANOVA with Šidák test. **** P < 0.0001, ns, no significant difference.

These findings suggest activity-dependent alterations in nAChR function at the NMJ in *nra-1* mutants. Specifically, NRA-1 may be required to sustain maximal efficiency of the levamisole receptor. To further test this hypothesis, we applied repeated 10 ms light stimuli at 0.5 Hz (Figure 3E-F). These experiments revealed a pronounced and persistent reduction in evoked currents in *nra-1* mutants (for comparison, see the same data normalized to the first peak current in Figure 3B, p < 0.0001). Together, these results demonstrate an activity-dependent requirement for NRA-1 in maintaining postsynaptic function of the cholinergic neuromuscular junction.

To further elucidate the molecular role of NRA-1 at L-AChRs and determine whether its loss affects receptor function, we recorded single-channel currents activated by ACh or levamisole in the cell-attached configuration from wild type and *nra-1(ok1025)* L1 muscle cells. The single-channel properties of wild type L-AChRs in *C. elegans* L1 muscle cells have been extensively characterized ^47,48^ and are consistent with those reported here.

Recordings of native channel activity elicited by 1 µM ACh from both wild type and *nra-1(ok1025)* L1 muscle cells revealed single-channel openings with comparable amplitudes (wild type: 3.5 ± 0.8 pA, n = 5; *nra-1(ok1025)*: 3.5 ± 0.9 pA, n = 5) at a pipette potential of 100 mV. Consistent with this, current–voltage analysis showed similar inward single-channel conductance for native L-AChRs in the presence or absence of NRA-1 (wild type: 35 ± 1.2 pS, n = 5; *nra-1(ok1025)*: 35 ± 2 pS, n = 5; p > 0.8), indicating that loss of NRA-1 does not affect the unitary conductance of the receptor (Figure 4A).

**Figure 4.**
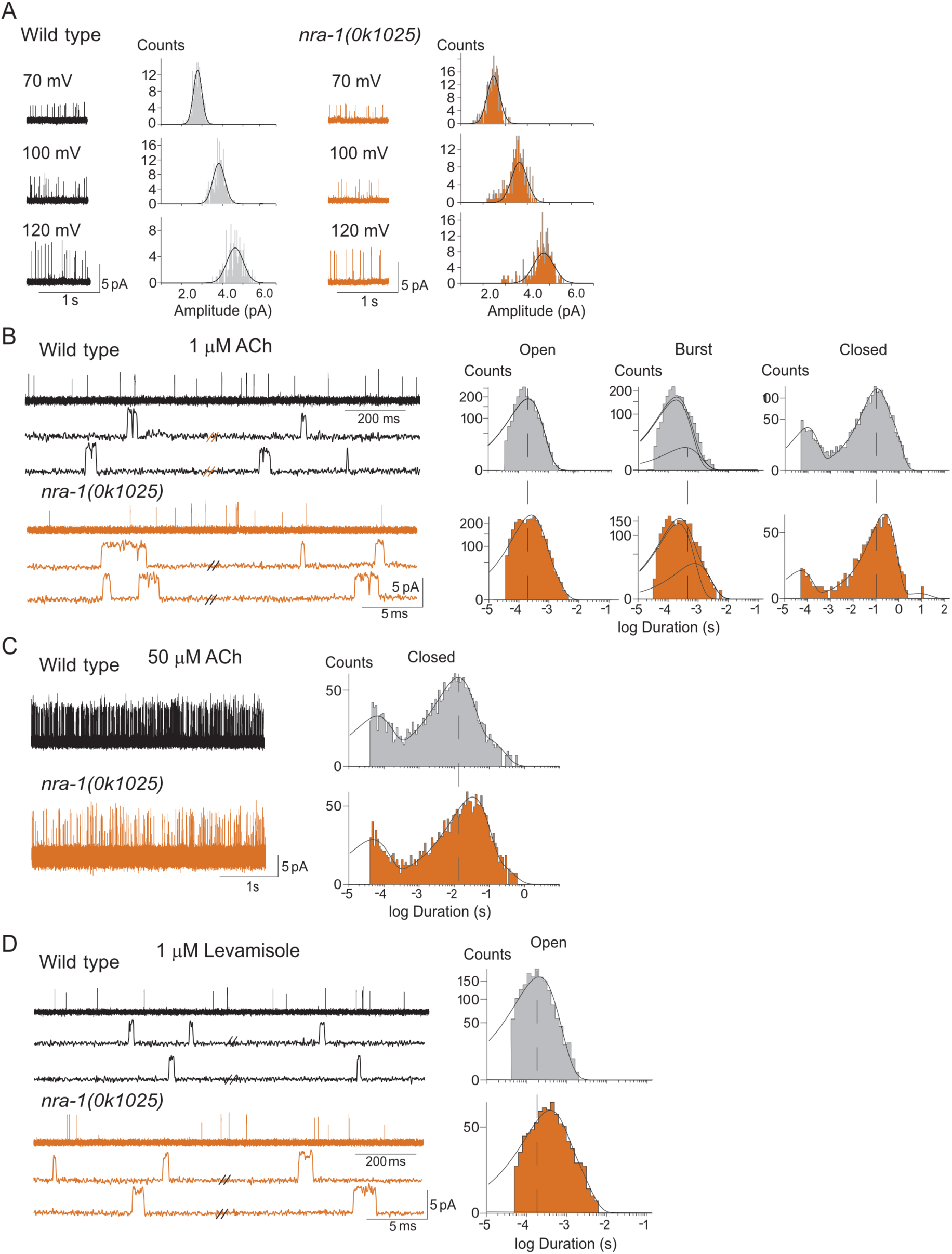
NRA-1 loss reshapes L-AChR single-channel gating without affecting unitary current amplitude. Single-channel recordings of native L-AChRs were obtained from primary cultures of L1 muscle cells isolated from wild type and *nra-1(ok1025) C. elegans* using the cell-attached configuration. Unless otherwise indicated, recordings were performed at a commanded pipette potential of +100 mV, filtered at 9 kHz, and channel openings are shown as upward deflections. A. Loss of NRA-1 does not alter L-AChR single-channel current amplitude. Left: Representative single-channel recordings activated by 1 µM ACh at the indicated pipette potential. Right: Corresponding amplitude histograms illustrating comparable unitary current amplitudes between wild type and *nra-1(ok1025)* L-AChRs. B. Loss of NRA-1 prolongs open and burst durations of ACh-activated L-AChRs. Left: Representative single-channel recordings evoked by 1 µM ACh, shown at two different time scales. Right: Open- Burst- and Closed-time duration histograms illustrating altered gating kinetics in *nra-1(ok1025)* L-AChRs. C. Reduced L-AChR activity at higher ACh concentrations in the absence of NRA-1. Representative single-channel recordings evoked by 50 µM ACh are shown for each strain. The corresponding closed-time duration histograms for each condition are also shown. D. Representative single-channel recordings activated by 1 µM levamisole are shown together with the corresponding open-time duration histograms for each strain.

In wild type muscle cells, open-time distributions of L-AChRs activated by 1 µM ACh were best described by the sum of two exponential components, comprising a predominant fast component (relative area > 0.80) with a mean duration of 0.24 ± 0.06 ms and a minor slow component of 0.48 ± 0.19 ms (Table 1). L-AChR in *nra-1(ok1025)* animals exhibited kinetic changes characterised by a significant increase in the duration of the slow open component (0.76 ± 0.14 ms; p < 0.05), while the predominant fast component remained comparable to that of wild type receptors in both duration and area (Figure 4B; Table 1).

**Table 1.** Single-channel properties of L-AChRs activated by ACh or levamisole in cultured C. elegans L1 muscle cells. Single-channel recordings were obtained from primary cultures of L1 muscle cells isolated from wild type and nra-1(ok1025) worms using the cell-attached configuration at a commanded pipette potential of +100 mV. The mean durations and relative areas of open-time components (O1 and O2) and the main closed-time components (C1 and C2) were derived from exponential fits to the corresponding dwell-time histograms. Data are expressed as mean ± SD, and n indicates the number of independent recordings analysed for each condition.

| Agonist | O1 (ms)<br>(area) | O2 (ms)<br>(area) | C1 (ms)<br>(area) | C2 (ms)<br>(area) | n |
| --- | --- | --- | --- | --- | --- |
| **Wild-type** |  |  |  |  |  |
| ACh |  |  |  |  |  |
| 0.1 μM | 0.23 ± 0.06<br>(0.82 ± 0.09) | 0.49 ± 0.10<br>(0.18 ± 0.12) | 0.08 ± 0.02<br>(0.16 ± 0.08) | 1508 ± 1093<br>(0.88 ± 0.12) | 5 |
| 1 μM | 0.24 ± 0.06<br>(0.48 ± 0.19) | 0.48 ± 0.19<br>(0.52 ± 0.16) | 0.06 ± 0.01<br>(0.12 ± 0.05) | 268 ± 232<br>(0.84 ± 0.11) | 5 |
| 50 μM | 0.22 ± 0.04<br>(0.45 ± 0.14) | — | 0.05 ± 0.01<br>(0.22 ± 0.02) | 15.0 ± 7.0<br>(0.70 ± 0.09) | 5 |
| Levamisole |  |  |  |  |  |
| 1 μM | 0.237 ± 0.047<br>(1.00) | — | 0.113 ± 0.059<br>(0.046 ± 0.015) | 99.4 ± 77.9<br>(0.88 ± 0.07) | 6 |
| ***nra-1(ok1025)*** |  |  |  |  |  |
| ACh |  |  |  |  |  |
| 0.1 μM | 0.29 ± 0.06<br>(0.80 ± 0.16) | 0.86 ± 0.20<br>(0.20 ± 0.16) | 0.045 ± 0.005<br>(0.22 ± 0.22) | 1247 ± 631<br>(0.75 ± 0.23) | 5 |
| 1 μM | 0.29 ± 0.10<br>(0.80 ± 0.04) | 0.76 ± 0.14<br>(0.26 ± 0.11) | 0.048 ± 0.007<br>(0.16 ± 0.04) | 640 ± 273<br>(0.80 ± 0.05) | 5 |
| 50 μM | 0.21 ± 0.03<br>(1.00) | — | 0.053 ± 0.002<br>(0.17 ± 0.03) | 38 ± 20<br>(0.83 ± 0.02) | 5 |
| Levamisole |  |  |  |  |  |
| 1 μM | 0.30 ± 0.07<br>(0.87 ± 0.06) | 0.91 ± 0.13<br>(0.13 ± 0.06) | 0.059 ± 0.027<br>(0.10 ± 0.03) | 493 ± 289<br>(0.83 ± 0.07) | 5 |

Analysis of burst durations, reflecting activation episodes of individual receptor molecules, revealed significantly longer mean durations in *nra-1(ok1025)* compared to wild type receptors (wild type: 0.64 ± 0.04 ms; *nra-1(ok1025)*: 1.1 ± 0.38 ms; p < 0.05) (Figure 4B), supporting a role of NRA-1 in shaping channel kinetics. We further analysed recordings obtained at lower ACh concentrations (0.1 µM), where channel activity is sparse. Open-time histograms from *nra-1(ok1025)* cells were best fitted by two exponential components, with the slowest component exhibiting a significantly longer duration than that observed in wild type cells (p < 0.01; Table 1), consistent with the effects observed at 1 µM ACh. Closed-time distributions recorded at 1 µM ACh in both wild type and *nra-1(ok1025)* muscle cells were best described by the sum of three to four exponential components. Notably, the predominant closed-time component, of approximately 300 ms in wild type cells, was significantly prolonged in *nra-1(ok1025)* mutants (about 600 ms; p < 0.05; Figure 4B; Table 1).

The prolongation of the predominant closed component may reflect a reduced number of functional receptors at the cell surface and/or altered channel kinetics associated with decreased channel opening. To directly assess this, we quantified the number of openings per unit time. Because the frequency of channel openings is concentration-dependent, measurements were obtained at 1 and 50 µM. A clear genotype-dependent difference was observed, with *nra-1(ok1025)* recordings exhibiting a markedly reduced opening frequency relative to wild type (Figure 4C). In wild type cells, opening frequencies were 11 ± 3 events/s at 1 µM ACh and increased to 100 ± 24 events/s at 50 µM ACh. In contrast, *nra-1(ok1025)* cells displayed significantly lower opening frequencies under both conditions, with values of 3 ± 1.2 events/s at 1 µM ACh (p < 0.001) and 36 ± 29 events/s at 50 µM ACh (p < 0.01; Figure 4C). Consistent with the reduced frequency of channel openings at 1 µM ACh, the predominant closed-time component at 50 µM ACh in *nra-1(ok1025)* recordings remained shifted toward longer durations (p < 0.05; Figure 4C), resulting in a significantly lower number of opening events per second. Together, these results indicate that loss of NRA-1 reduces the overall activity of L-AChRs.

To extend the analysis, we recorded single-channel activity evoked by 1 µM levamisole in wild type and *nra-1(ok1025)* L1 muscle cells. Recordings at 1 µM levamisole revealed clear qualitative differences between wild type and *nra-1(ok1025)* muscle cells. At this concentration, open-time histograms from wild type L-AChRs were best described by a single exponential component. In contrast, recordings from *nra-1(ok1025)* cells required two exponential components to adequately describe the open-time distributions, revealing the presence of a longer open component that was absent in wild type receptors (0.9 ± 0.4 ms, Figure 4D and Table 1). Consistent with the effects observed for ACh-activated channels, closed-time distributions at 1 µM levamisole also showed a marked shift of the predominant closed component toward longer durations in *nra-1(ok1025)* cells compared to wild type, contributing to an overall reduction in receptor activity (p < 0.01; Table 1).

Guided by our findings that loss of NRA-1 results in a clear reduction in optogenetically evoked ACh currents, as well as an overall reduced activity of the L-AChR, we next sought to determine how loss of *nra-1* affects behaviour in the worm. To this end, we first assessed basic locomotion function off food in the *nra-1(ok1025)* strain and wild type N2 worms and could conclude that *nra-1(ok1025)* is not defective in any basic locomotor parameter examined (total speed, forward and backward speed, reversal frequency and speed) (Figure S1). We next investigated sensory function by assessing the chemotactic response of *nra-1(ok1025)* mutants to the well-established attractant odours diacetyl and isoamyl alcohol (IAA) (Figure 5A, C). We found that *nra-1(ok1025)* animals were significantly impaired in chemotaxis towards both diacetyl and IAA compared to wild type (CI_diacetyl_ wild type: 0.85 ± 0.02, *nra-1(ok1025):* 0.62 ± 0.03, p < 0.0001, CI_IAA_ wild type: 0.84 ± 0.02, *nra-1(ok1025)*: 0.66 ± 0.04, p < 0.001;Figure 5B, D).

**Figure 5.**
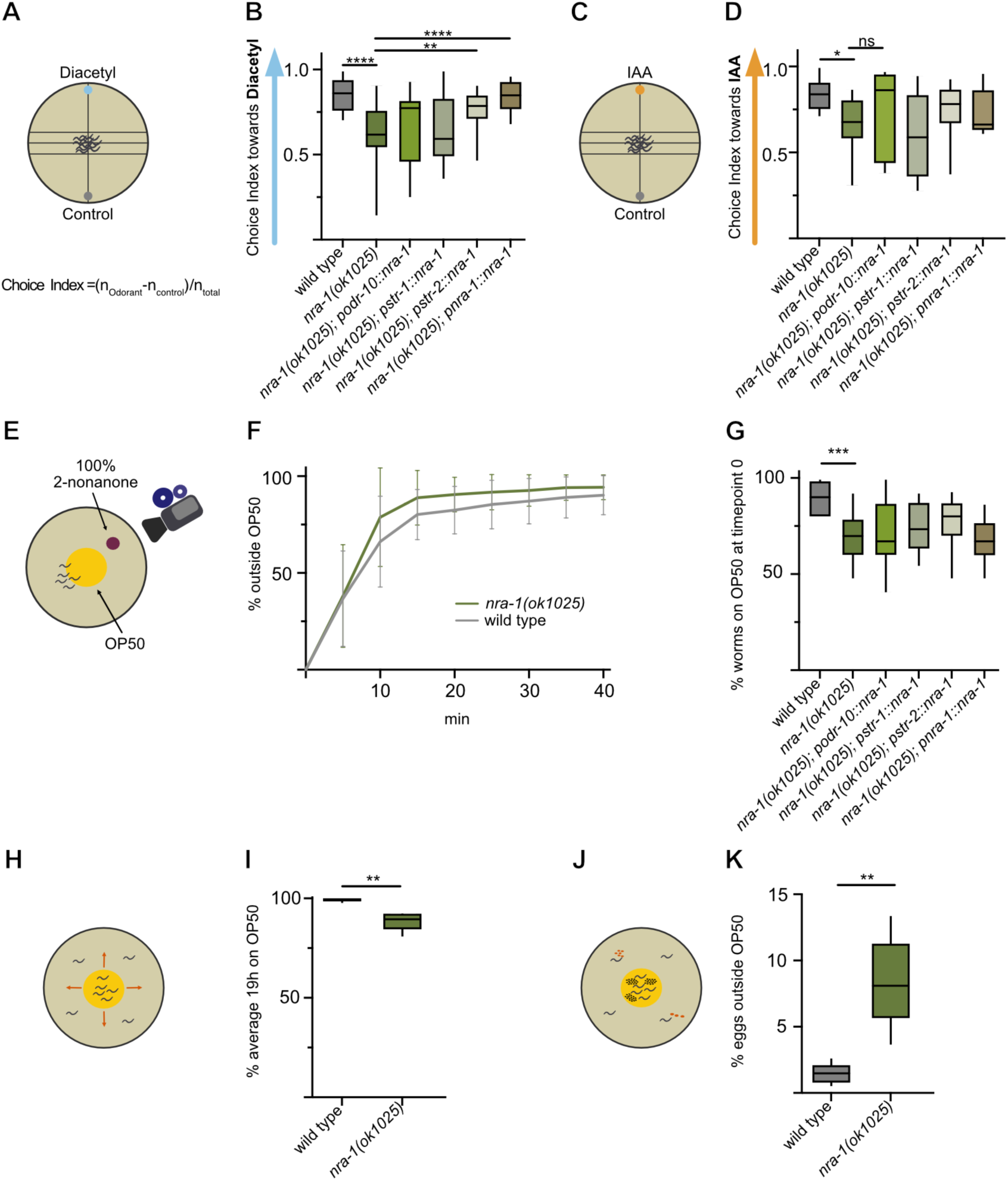
*nra-1* mutants are deficient in their chemotactic behaviour towards both diacetyl and IAA with abnormal food-sensing behaviour. A, C. Description of chemotaxis experiment. B, D. *nra-1(ok1025)* mutants are defective in chemotaxis towards diacetyl and IAA. The phenotype can be rescued for diacetyl by expression of *nra-1* in AWC (*pstr-2::nra-1*) and under its own endogenous promotor. E. Experimental setup for 2-nonanone food leaving experiment. F. Food-leaving in the presence of 100% 2-nonanone. G. Percentage of worms found on the food patch at timepoint zero, only the wild type and *nra-1(ok1025)* worms showed a significant difference at timepoint 0. H. Experimental outline of long-term food-leaving experiment. I. Percentage of worms on the food patch after 19h. J. Experimental outline of egg-laying experiment. K. Percentage of eggs laid off the food patch. N=10-25 plates/genotype, *P < 0.05, **P < 0.01, ***P < 0.001, ns, no significant difference, calculated by Welch’s t-test, data are shown as mean ± SEM.

To verify that these phenotypes were due to the loss of *nra-1* function, we performed rescue experiments using different promotor constructs. Expression of *nra-1* under its endogenous promotor successfully restored chemotaxis towards diacetyl, but not towards IAA (CI_diacetyl_ *nra-1(ok1025);pnra-1::nra-1*: 0.84 ± 0.03, p < 0.005, CI_IAA_: 0.76 ± 0.3, p = 0.15; Figure 5B, D). Interestingly, the diacetyl chemotaxis defect was also rescued when *nra-1* was expressed in the AWC^ON^ neuron using the *str-2* promotor (CI_diacetyl_ *nra-1(ok1025);pstr-2::nra-1*: 0.76 ± 0.03, p < 0.05; Figure 5B), but not by using the AWA-specific *odr-10* promotor (CI_diacetyl_ *nra-1(ok1025);podr-10::nra-1*: 0.67 ± 0.05, p = 0.45; Figure 5B). The ability of AWC^ON^-specific *nra-1* expression to restore chemotaxis to diacetyl, even though the diacetyl sensing receptor ODR-10 is not expressed in these neurons, suggests that NRA-1 might act beyond the level of primary odour detection. Instead, NRA-1 may function at a circuit-integration or modulatory level, where it could influence neuronal communication or synaptic strength between sensory neurons such as AWA (the primary diacetyl-sensing neuron^12^) and interneurons downstream of AWC. In addition, we were not able to rescue the chemotactic behaviour towards IAA by re-introducing *nra-1* into the AWC neuron, the neuron that senses IAA^64^ (CI_IAA_ *nra-1(ok1025);pstr-1::nra-1*: 0.60 ± 0.07, p = 0.52; Figure 5D). This finding supports a model in which NRA-1 contributes to the coordination of sensory input across neuronal subtypes, promoting appropriate behavioural output in response to environmental cues.

To further investigate whether the *nra-1* mutant exhibits additional sensory deficits, we tested its ability to detect the repulsive odour 2-nonanone in the presence of food. 2-nonanone is sensed by the AWB neurons, and wild type *C. elegans* are strongly repelled by it^15^. In contrast, under normal conditions worms leave a bacterial food source only at low frequency^65^. We therefore examined whether exposure to 2-nonanone in the presence of food would influence the worms’ food-leaving behaviour (Figure 5E). After 10 minutes, food-leaving was pronounced in both genotypes, with most worms having left the food patch (Figure 5F). Although no significant differences were detected between the strains, it was noted that at the start of the experiment (timepoint zero), a substantial fraction of *nra-1* mutants was already positioned outside the food patch, suggescng that they had le{ the lawn during the one-hour recovery period prior to the assay (% on food at t=0: wild type: 89.4 ± 2.37, *nra-1(ok1025):* 68.9 ± 4.1, p < 0.001; Figure 5G). To determine whether this was a transient effect caused by transferring the worms onto a new plate, we examined their distribution after 19 hours. At this time point nearly all wild type worms remained on the food, whereas a significantly lower proportion of *nra-1* mutants were found within the food patch (% on food at t=19h: wild type: 99 ± 0.44, *nra-1(ok1025):* 88.3 ± 1.93, p < 0.01; Figure 5I). To further assess this phenotype, we quantified egg-laying behaviour relative to the food source (Figure 5J). Wild type worms typically retain their eggs when away from food^66^, and defects in sensory processing, such as dysfunction in the dopaminergic PDE neurons, strongly affect this behaviour^66–68^. Consistent with a sensory impairment, *nra-1* mutants laid a significantly higher proportion of eggs off the food compared to wild type worms (% eggs outside food t=19h: wild type: 1.47 ± 0.34, *nra-1(ok1025):* 8.35 ± 1.43, p < 0.01; Figure 5K), indicating an inability to correctly perceive the boundaries of the food source. Together, these behavioural results suggest that *nra-1* mutants exhibit a global sensory deficit that impairs their capacity to sense and localise food, leading to abnormal food-seeking and egg-laying behaviours without a direct impairment of NMJ function and basal locomotion.

## Discussion

Here, by combining electrophysiological and behavioural analyses, we demonstrate that NRA-1 is required to maintain efficient postsynaptic cholinergic receptor function in an activity-dependent manner and contributes to sensory processing. The absence of detectable locomotor defects argues against a general impairment of motor output and instead suggests that loss of NRA-1 primarily affects neuronal circuits involved in sustained synaptic activity, sensory integration, and behavioural state transitions. By linking NRA-1 function across both single-receptor and whole-organism scales, our findings extend the functional repertoire of copine proteins in the nervous system and highlight their conserved roles across species.

Our initial observations showed that acute agonist application evoked comparable macroscopic currents in wild type and *nra-1(ok1025)* mutants. This finding was unexpected given our previous finding of reduced L-AChR abundance at the muscle membrane^26^ and suggested that overall receptor expression, or function, is not grossly impaired under non-physiological stimulation conditions. Because pressure application activates receptors across the entire muscle surface, this assay is likely to obscure defects in synaptic receptor positioning or re-positioning. Indeed, when synaptic transmission was probed using optogenetic stimulation of cholinergic neurons, clear genotype-dependent differences emerged. While initial responses to brief or prolonged stimulation were largely preserved, sustained responses and repeated stimulation paradigms revealed a pronounced and progressive reduction in evoked currents in *nra-1* mutants. These findings indicate that NRA-1 is not required for baseline receptor function, but rather for maintaining efficacy in NMJ signalling during ongoing or repeated activity. This interpretation is consistent with literature showing that postsynaptic receptor trafficking and nanoscale localisation can strongly influence synaptic efficacy even when total receptor abundance or agonist-evoked responses appear relatively preserved^30,69^.

This activity-dependent phenotype suggests a role for NRA-1 in stabilising postsynaptic receptor function, potentially through regulating receptor localisation or retention at synaptic sites. Such a mechanism is consistent with the known properties of copine proteins as calcium-dependent membrane-binding adaptors^27^, which can couple signalling events to membrane trafficking processes. In this context, NRA-1 may facilitate the recruitment or maintenance of L-AChRs at postsynaptic domains during periods of elevated synaptic activity, thereby sustaining efficient neurotransmission.

Our single-channel analyses provide mechanistic insight into how loss of NRA-1 alters receptor function. Importantly, unitary conductance was unchanged in *nra-1(ok1025)* mutants, indicating that NRA-1 does not affect the fundamental ion-permeation properties of L-AChRs. Instead, NRA-1 selectively modulates channel activity. In the absence of NRA-1, L-AChRs exhibited prolonged open and burst durations, together with a significant increase in closed-state durations and a marked reduction in the frequency of channel openings. These alterations were consistently observed across agonist concentrations and were reproduced with both ACh and levamisole, indicating that NRA-1 broadly regulates receptor activity rather than mediating ligand-specific effects. The prolonged closed states and reduced opening frequency observed in the absence of NRA-1 reflect diminished agonist-evoked channel activity. This reduction may arise because NRA-1 promotes efficient channel activation or helps maintain an adequate number of functional receptors. The changes in open and burst durations further support a role for NRA-1 in modulating channel-gating kinetics. Together, these findings indicate that NRA-1 is required to ensure normal L-AChR function and, consequently, proper neuromuscular transmission.

An important question raised by our findings is how the activity-dependent defects observed at the cholinergic NMJ relate to the pronounced sensory phenotypes of *nra-1* mutants. Although loss of *nra-1* resulted in reduced sustained cholinergic responses during repeated stimulation, mutants did not display detectable defects in baseline locomotion, suggesting that the behavioural abnormalities cannot be explained solely by impaired motor output. Instead, the rescue of diacetyl chemotaxis by AWC^ON^-specific expression of *nra-1,* despite these neurons not being primary detectors of this odour^12^, points to a role within sensory integration circuits. Notably, many interneurons that relay chemosensory information in *C. elegans* are cholinergic, raising the possibility that the activity-dependent cholinergic defects observed at the NMJ reflect a broader requirement for NRA-1 at cholinergic synapses throughout the nervous system. In this model, the NMJ serves as a tractable system in which NRA-1-dependent regulation of receptor function can be studied directly, while similar mechanisms may influence information processing in sensory circuits. This interpretation is consistent with the broad neuronal and muscle expression pattern of *nra-1* and with the organisation of *C. elegans* sensory circuits, in which behavioural output depends on integration across multiple sensory neurons, interneurons, premotor neurons and followed by neuromuscular activation, rather than on primary odour detection alone^38,70^. An alternative, non-exclusive possibility is that NRA-1 regulates additional ligand-gated ion channels beyond L-AChRs, including receptors operating at glutamatergic sensory-interneuron synapses. Future studies will be required to determine the extent to which NRA-1 acts as a general regulator of ligand-gated ion channel function across different neuronal cell types.

While mammalian copines have been implicated in synaptic plasticity and membrane protein localisation^27,28,30,71^, their roles in intact neural circuits have remained poorly understood. Our results provide *in vivo* evidence that a copine protein can regulate ion channel function in an activity-dependent manner, linking membrane trafficking to synaptic efficacy and behaviour. This suggests that copine-mediated regulation of receptor dynamics may represent a conserved mechanism for tuning neuronal responsiveness across species.

In summary, we propose that the copine protein NRA-1 acts as an activity-dependent regulator of postsynaptic cholinergic receptor function, promoting efficient channel gating and sustaining synaptic transmission during repeated activity. Through this role, NRA-1 supports functions in sensory circuits and enables appropriate behavioural responses to environmental cues. These findings establish a direct connection between membrane-associated regulatory proteins, ion channel function and sensory behaviour that opens new avenues for understanding how receptor trafficking and kinetics are coordinated in nervous systems.

## Acknowledgements

The authors gratefully acknowledge past and present members of the Morud Lekholm lab for helpful discussions. This work was supported by grants from the Swedish Research Council (2022-03951), Swedish Foundation for Strategic Research (FFL21-0166), Tore Nilssons stiftelse, Åke Wibergs stiftelse (M22-0215, M21-0229), Magnus Bergvalls Stiftelse (J.M), Lundgrenska stiftelserna (L.C), Deutsche Forschungsgemeinschaft (grant ID GO1011/3-2 and EXC115 to AG), Argentine grants from Universidad Nacional del Sur (PGI 24/B366 to GH and PGI 24/B360 to CB), Agencia Nacional de Promoción de la Investigación, el Desarrollo Tecnológico y la Innovación (PICT 2020 − 00936 to CB) and Consejo Nacional de Investigaciones Científicas y Técnicas (CONICET, PIP11220200102356).

## Author contribution

Conceptualisation: J.M., A.G., C.B., L.C., data curation: L.C., J.F.L., E.A., G.H., methodology: J.M., L.C., A.G., C.B., J.F.L., S.W., G.H., investigation: L.C., E.A., J.F.L., S.W., G.H., formal analysis: L.C., J.M., C.B., A.G., J.F.L., G.H., visualisation: L.C., J.M., funding acquisition: J.M., A.G., C.B., project administration: J.M., supervision: J.M., A.G., C.B., validation: J.M., A.G., C.B., resources: J.M., A.G., C.B., writing – original draft: J.M., A.G., C.B., writing – review and editing: J.M., A.G., C.B., L.C., E.A.

## Data and code availability

All experimental data, including results of imaging, behavioural and biochemical experiments, will be shared by the lead contact upon request. Any additional information required to re-analyse the data reported in this paper is available from the lead contact upon request.

## Supporting Figures

**Supporting Figure 1:**
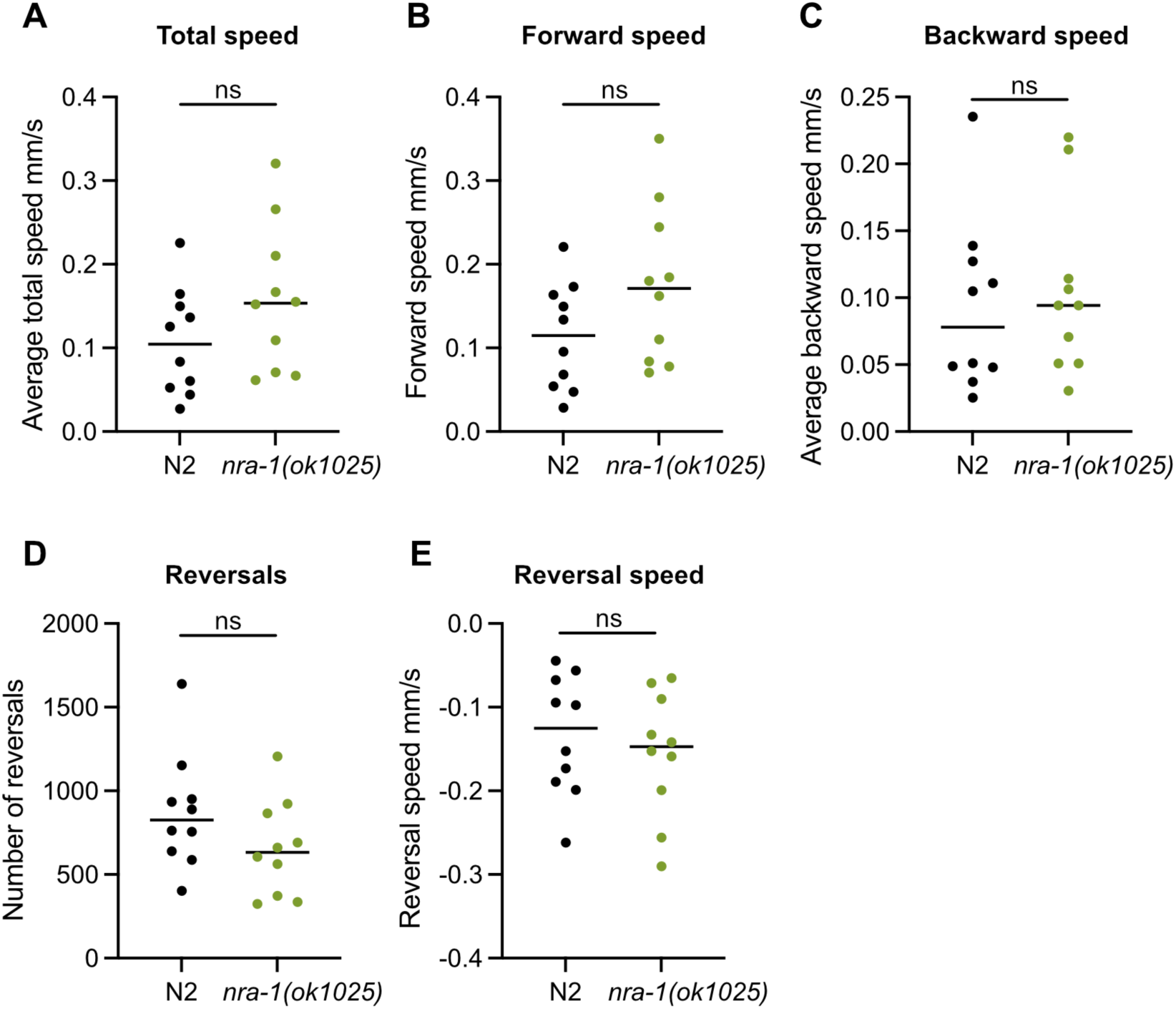
*nra-1* mutants are not defective in any basic locomotor function tested as compared to N2 wild type worms. All tracking was performed off food. A. Average total locomotion speed in mm/s. B. Forward locomotion speed in mm/s. C. Average backward locomotion speed in mm/s. D. Number of reversal movements detected during a 10-minute experiment. E. Average reversal speed during reversal behaviour in mm/s. N=10 plates/genotype with 12-16 worms/plate, ns, no significant difference, calculated by Welch’s t-test, data are shown with a scatter dot plot, the line represent the median value.

